# Temporal physiological, transcriptomic and metabolomic analyses revealed molecular mechanism of *Canna indica*’s response to Cr stress

**DOI:** 10.1101/2023.01.14.524062

**Authors:** Zhao Wei, Chen Zhongbing, Yang Xiuqing, Sheng Luying, Mao Huan, Zhu Sixi

## Abstract

Chromium (Cr) can interfere with plant gene expression, change the content of metabolites and affect plant growth. However, the molecular response mechanism of wetland plants at different time sequences under Cr stress has yet to be fully understood.The results showed that Cr stress increased the activities of superoxide dismutase (SOD), ascorbate peroxidase (APX) and peroxidase (POD), the contents of glutathione (GSH), malondialdehyde (MDA), and oxygen free radical (ROS), and inhibited the biosynthesis of photosynthetic pigments, thus leading to changes in plant growth and biomass. that Cr stress mainly affected 12 metabolic pathways, involving 38 differentially expressed metabolites, including amino acids, phenylpropane, and flavonoids. A total of 16247 differentially expressed genes were identified, among which, at the early stage of stress, *C. indica* responds to Cr toxicity mainly through galactose, starch and sucrose metabolism. With the extension of stress time, plant hormone signal transduction and MAPK signaling pathway in *C. indica* in the treatment group were significantly affected. Finally, in the late stage of stress, *C. indica* co-defuses Cr toxicity by activating its Glutathione metabolism and Phenylpropanoid biosynthesis. In conclusion, this study revealed the molecular response mechanism of *C. indica* to Cr stress at different times through multi-omics methods.

**Graphical Abstract:** 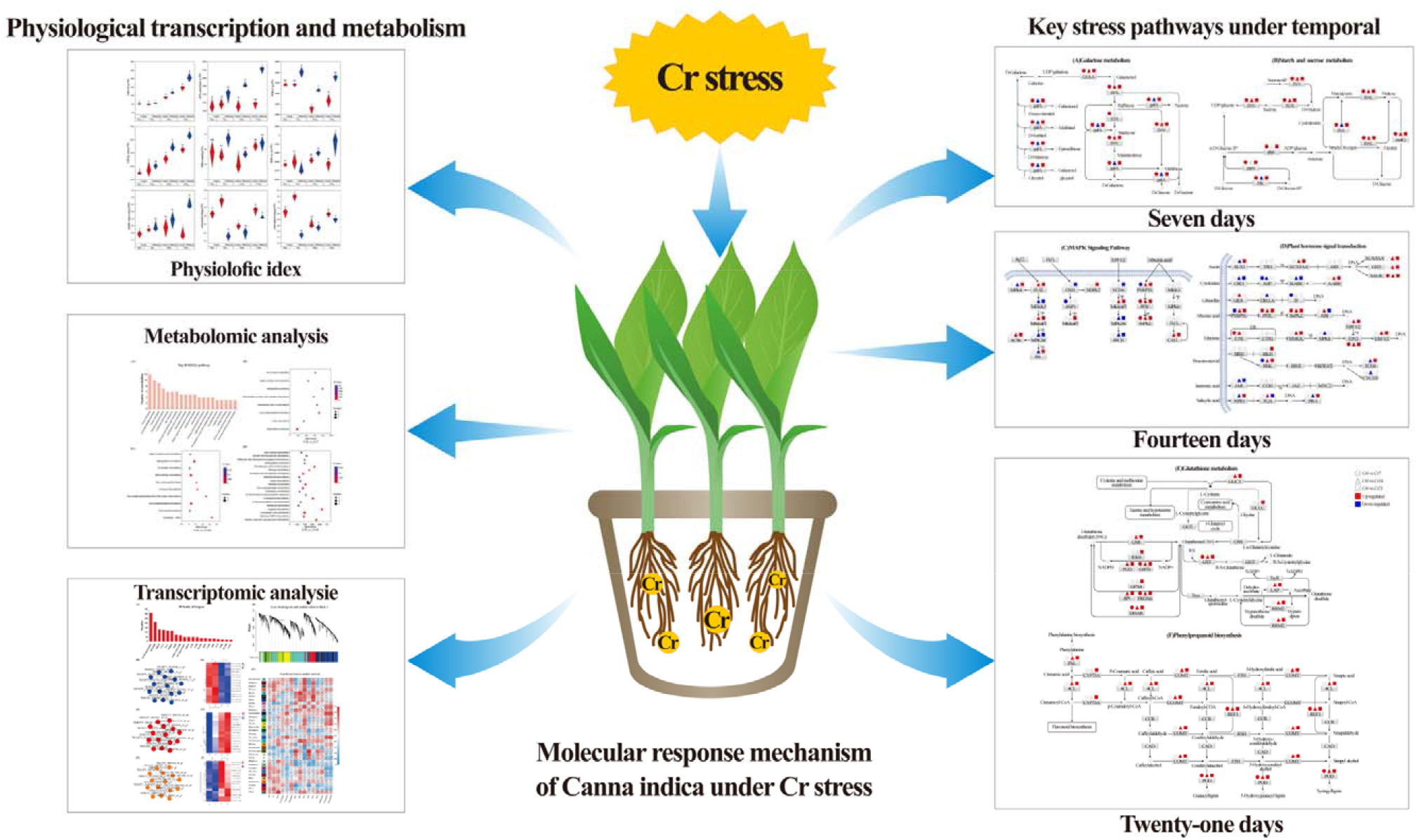

## 1. Introduction

Heavy metal pollution has become a global environmental problem (Uchimiya et al., 2020). In recent years, due to the influence of human activities, Cr has been widely distributed in soil and water (Xing et al., 2021). Among them, industrial activities (such as electroplating, smelting, and mining) and agricultural activities (such as pesticide use and fertilizer application) are the primary sources of Cr pollution (Hakan et al., 2021). In nature, Cr exists in trivalent (Cr^3+^) and hexavalent (Cr^6+^) forms, with Cr^6+^ having good mobility and toxicity (Ahmad et al., 2020). Although Cr is a non-essential element in plants, it can still accumulate in large amounts in plant roots and aboveground parts (Yu et al., 2018). Even trace levels can harm plants’ morphological, physiological, and molecular characteristics (Arun et al., 2005). In addition, at high concentrations, Cr can lead to a variety of toxic symptoms in plants, such as inducing oxidative stress, resulting in excessive production of reactive oxygen species (ROS), reducing the activity of antioxidant enzymes, blocking the synthesis of photosynthetic pigments, inhibiting photosynthesis, thus affecting plant growth and development and reducing their biomass (Sinha et al., 2018). In addition, Cr is a non-biodegradable heavy metal element that can exist in plants for a long time and pose a potential threat to human and animal health through its spread through the food chain (Riti et al., 2022). Therefore, the remediation of Cr-contaminated soil is necessary and urgent.

In the past few decades, several remediation methods for Cr contamination have emerged, among which physical, chemical, and biological methods have been successfully applied to the remediation of Cr-contaminated soils (Singh et al., 2022). Compared with traditional restoration techniques, phytoremediation is an aesthetic, economical, and publicly recognized in-situ bioremediation technology (Ashraf et al., 2019). It can provide an effective solution for soil removal, transfer, degradation, and fixation of Cr (Ao et al., 2022). However, some challenges remain in its application, as phytoremediation mainly depends on the concentration of Cr in the soil (Anastasis et al., 2021). Therefore, screening Cr-tolerant plants and understanding the molecular response mechanism of these plants on Cr tolerance is the focus of phytoremediation research, which will further promote the remediation effect of Cr-contaminated soil (Vibha et al., 2018). At present, some Cr-tolerant plants have been identified by studies. There are *Leersia hexandra* (Zhang et al., 2022a), *C. indica* (Zhao et al., 2017), *Cyperus alternifolius* (Wang et al., 2021a), *Medicago sativa* (Wang et al., 2012). These plants have evolved various defense and detoxification mechanisms to cope with heavy metal chromium (HMC) stress. Among them, the cell wall is the first physical barrier that effectively inhibits Cr from entering root cells (Wang et al., 2021b; Yuan et al., 2022a), which can significantly reduce Cr absorption and fix Cr in the cell wall. When the first barrier is breached, plants activate their antioxidant defense system and use vacuoles for compartments, thus alleviating the toxic effects of Cr (Zhong et al., 2019). However, due to the diversity of plants, we need to explore whether there is a common response mechanism among different Cr-accumulating plants.

*C. indica* belongs to Canna indica, a perennial herbaceous species, showing a developed root system, strong adaptability to the living environment, rapid growth, large leaf area and other characteristics, and strong enrichment ability to heavy metals (Zhang et al., 2012). In treating contaminated wastewater containing heavy metals such as Cr, it is found that it has a robust, comprehensive tolerance, which can quickly adjust its own physiological and biochemical characteristics, showing strong tolerance (Liu et al., 2011). At present, a large number of studies have focused on the response mechanism of *C. indica* to Cr stress in morphology, physiology, and biochemistry, including plant biomass, plant chelate (PC) synthesis, photosynthetic pigment, antioxidant defense system, and organic acid secretion (Dong et al., 2019; Zhang et al., 2020; Xiang et al., 2022). Our previous studies showed that the contents of chlorophyll, malondialdehyde (MDA), and reduced glutamate (GSH) in *C. indica* seedlings changed significantly with increased Cr concentration. Moreover, the activities of enzymes related to the antioxidant mechanism (SOD, CAT, POD, and PAL) were also changed (Zhao et al., 2017). However, the underlying molecular mechanism of *C. indica* under Cr stress remains largely unknown. In recent years, with the development of transcriptome sequencing (RNA-Seq) and metabolomics techniques, they have been widely used to reveal the different response mechanisms of different plants to Cr stress. Including *Zea mays* (Hakan et al., 2021), *Helianthus annuus* (Ibarra et al., 2019), *Arabidopsis thaliana* (Jia et al., 2016), *Sorghum bicolor* (Roy et al., 2016). Therefore, by taking *C. indica* as the research object and combining it with multi-omics techniques, we can improve the knowledge gap of *C. indica’s* response to Cr stress at the molecular level.

Therefore, physiological, transcriptomic, and metabolomic methods were used in this study to analyze the molecular response mechanism of *C. indica* to Cr stress at different exposure times. We hypothesized that Cr stress could induce the abnormal expression of many genes and metabolites related to the antioxidant and detoxification mechanisms of *C. indica*. The purpose of this study was to: (a) analyze the accumulation and transport of Cr by *C. indica* and its physiological changes under Cr stress at different exposure times; (b) identify their key metabolic pathways in differential expressed genes (DEGs) and differentially expressed metabolites (DEMs); (c) reveal the molecular mechanism of *C. indica* tolerance under Cr stress at different exposure times, to provide a theoretical basis for phytoremediation of soil Cr pollution, and identify essential phytoremediation candidate genes to provide a theoretical basis for future research.

## 2. Materials and methods

### 2.1. Plant cultures and Cr treatment

The *C. indica* seedlings used in this study were hydroponically grown by Songnan Plant Seedling Company of Luzhi town, Suzhou City. Firstly, *C. indica* seeds of the same size were screened. After sterilization, seeds were cultured in Petri dishes, waiting for germination, and then moved to nutrient water for hydroponics. When the plants grew to about 10 cm, we purchased seedlings from the company and selected seedlings with similar growth conditions for the experiment. The selected *C. indica* seedlings were surface disinfected with 75% ethanol and 1% sodium hypochlorite solution for 10 s and 15 min. Carefully washed with deionized water five times, furthermore transplanted in the potted greenhouse (500g of soil per pot). All seedlings were then earth culture in a controlled greenhouse (176 μmol m^2^ s^-1^ light intensity, 12 h photoperiod, 25 °C constant temperature). Seedlings were domesticated in the greenhouse for 30 days before exposure to Cr. Hoagland solution (15 mL) and water (15 mL) were added to the pot every 15 days and every three days, respectively, and Hoagland nutrient solution was not added after Cr stress. Twenty-one seedlings with similar growth conditions were randomly divided into seven groups with three plants in each group (as three biological replicates): four groups were the control group (0 mg/kg K_2_Cr_2_O_4_), and the other three groups were the Cr treatment group (100 mg/kg K_2_Cr_2_O_4_). Seedlings were treated for 0, 7, 14, and 21 days before harvest (three plants at 0 days, six plants at 7 days, six plants at 14 days, and six plants at 21 days). Root (R) tissue from both Cr-treated and untreated seedlings was sampled simultaneously on corresponding days and immediately frozen in liquid nitrogen and stored in a −80 °C freezer until further treatment. Finally, 21 samples were collected and prepared for transcriptomic and metabolomic analysis. Similarly, samples (21 seedlings) for biochemical analysis were collected in this manner.

### 2.2. Cr content in soil and plants

After the harvest of plant samples, all soil samples were placed on yellow paper and air-dried at the vent. After air drying, soil samples were repeatedly knocked in a cloth bag, screened with 100 mesh, and then placed in a separate sealed bag for subsequent determination. After weighing 0.5 g of soil sample and preparing to reverse king’s water with a ratio of 1:3 (concentrated hydrochloric acid and concentrated nitric acid), the two were added into the digestion tube in turn, and the digestion was heated on the electric heating plate under the fume hood. First, the digestion was conducted at 160 °C for one hour, then 3 ml perchloric acid was added for another hour after the temperature was raised to 200 °C, and the remaining liquid was moved to the test tube. It was fixed with 1% dilute nitric acid. Plant samples (directly after harvest) were dried in a 105 °C oven for 12 hours. After that, the samples were ground to a diameter of less than 0.02 mm. The plant samples (0.5 g of each) were added with 37% HCl and 63% HNO_3_, sealed, and put into the oven at 150°C for digestion for 10 hours. Then dilute the suspension with 3 ml HNO_3_. The Cr analysis for soil and plant samples was performed using inductively coupled plasma mass spectrometry (ICP-MS) (Agilent, 7800 ICP-MS, USA) (Meng et al., 2022).

### 2.3. Soil indicator and Physiological index of plants

#### 2.3.1. Soil Physicochemical Properties

Soil pH and EC was measured by the potentiometric method. Soil organic matter (SOM) was measured by K_2_Cr_2_O_7_-H_2_SO_4_ oxidation-external heating method (Bao, 2000).

#### 2.3.2. Chlorophyll and carotenoid contents

Two leaves of 0.1g *C. indica* were collected and placed in a mortar with a small amount of powder (about 50 mg) and a small amount of Chlorophyll Assay Buffer and Carotenoid Assay Buffer, respectively. Then it was ground into homogenate, transferred to a 10 ml centrifuge tube, supplemented with leaves of Chlorophyll Assay Buffer and Carotenoid Assay Buffer to 10 ml, and placed away from light for 5 min-2 h. When the tissue was close to white and pigment extraction was completed, after centrifugation, the supernatant was taken to determine chlorophyll a, chlorophyll b, and carotenoids at 665 nm, 649 nm, and 470 nm, respectively (Fan et al., 2018).

#### 2.3.3. SOD, POD, and APX contents

Superoxide dismutase (SOD, EC 1.15.1.1) and peroxidase (POD, EC 1.11.1.7) in *C. indica* leaves were determined using a specified kit according to a protocol provided by manufacturer Suzhou Keming Biotechnology Co., LTD (www.cominbio.com). and ascorbate enzyme (APX) levels. Conditions 5 (BDTS, USA) multifunctional ELISA measured absorbance at 560 nm, 240 nm, 470 nm, and 290 nm, respectively.

#### 2.3.4. MDA, GSH and ROS levels in plants

For MDA analysis, 10% cold trichloroacetic acid was added into 0.1g ground sample and centrifuge at 4000 r/min for 10 min. Then, 2 ml 0.6% thiobarbituric acid was added to the samples and treated in a 100 °C water bath for 15 min. The supernatant was quickly cooled and centrifuged again. Absorbance was measured at 600 nm and 532 nm, and MDA content was calculated following the previous description (Mbonankira et al., 2015).

The aerial parts of 0.1g of *C. indica* were weighed, ground with EDTA-TCA reagent, and diluted in a 25 ml volumetric flask. Then, 2 ml of the filtrate was added to 0.4 ml of 1 mol/L NaOH reagent, and PBS reagent and 0.1 ml of TDNB reagent were added to the solution at pH 6.5-7.0; the test tube to which only K_3_PO_4_ was added served as a control. Finally, the samples were incubated at 25°C for 5 min to allow full-color development, absorbance was measured at 412 nm, and GSH content was calculated following the previous description (Meng et al., 2022).

Reactive oxygen species (ROS) were detected by fluorescent probe DCFH-DA using a specified kit. Dcfh-da itself has no fluorescence and can freely cross the cell membrane. After entering the cell, it can be hydrolyzed by intracellular esterase to produce DCFH. However, DCFH cannot penetrate the cell membrane, making it easy for the probe to be loaded into the cell. Moreover, intracellular reactive oxygen species can oxidize non-fluorescent DCFH to produce fluorescent DCF. Detecting the fluorescence of DCF can measure the level of Reactive oxygen species (ROS) in the leaves of *C. indica*. Conditions 5 (Berten, USA) multifunctional ELISA instrument was used to measure the fluorescence intensity for 10 min, with an excitation wavelength of 499 nm and emission wavelength of 521 nm.

#### 2.3.5. Soluble matter contents in plants

Soluble sugar content was measured following the guidelines of Gao et al. (Gao et al., 2006). First, 0.1g leaves were placed in a test tube filled with distilled water, boiled at 100 °C for 20 min, and cooled in a 100 ml volumetric bottle at constant volume. Then, 1 ml was absorbed, and 5 ml of anthrone-measuring reagent was added. After the mixture was treated in boiling water, absorbance was measured at 620 nm, and soluble sugar content was calculated.

### 2.4. Transcriptome analysis

Novaseq 6000 (Illumina) was used for transcriptome sequencing to understand further the molecular response mechanism of Cr detoxification in *C. indica* to determine the changes in gene expression in the roots of *C. indica* under Cr stress at different times. Root tissue (3 replicates per group) of 0.5 g was taken from Cr-treated and untreated *C. indica* seedlings at days 0, 5, 10, and 15 and were frozen in liquid nitrogen. The RNA of root samples of *C. indica* under CK and Cr treatments (including three biological replications) was extracted by TRIzol® Reagent (Invitrogen, USA), purified by Plant RNA Purification Reagent (Invitrogen company), and sequenced on the HiSeq 6000 Illumina sequencing platform by Shanghai Majorbio Bio-pharm Technology Co. Ltd, China. The DEGs of the root of *C. indica* between the CK and Cr treatments were identified by DEGseq2. The functional annotation of DEGs was subjected to Swiss-Prot annotation, Clusters of Orthologous Groups of proteins (COG), Gene Ontology (GO) classification, and Kyoto Encyclopedia of Genes and Genomes (KEGG) database by the free online platform of Majorbio (www.majorbio.com).

### 2.5. Metabolomics analysis

Root tissue of 0, 7, 14, and 21-day time series from Cr-treated and untreated *C. indica* seedlings (3 replicates per group) was taken at about 1 g (3 replicates per group), weighed, and frozen in liquid nitrogen. After natural air drying, root samples were ground to powder, and metabolites were extracted and analyzed. The amount of 60 mg ground powder was ultrasonically extracted with 0.6 ml methanol/water (7:3, v/v) for 30 min followed by 20 min incubation at −20 °C, coupled with an internal standard of L-2-Cl-Phe (0.3 mg ml^-1^). The extracts were centrifuged at 14,000 rpm for 10 min at 4 °C. Then, 200 μl of supernatant was filtered through a 0.2 μm filter and measured using a Waters VION IMS Q-TOF Mass Spectrometer equipped with an electrospray interface (Waters Corporation, Milford, MA, USA) platform as described elsewhere (Su et al., 2021).

### 2.6. Statistical analysis

Shapiro-Wilk and Levene’s tests were performed to test the normality and homogeneity of data. The natural logarithm was applied to transform the data that did not obey a normal distribution. Statistical analyses were performed by SPSS software (version 26.0). All values are expressed as the mean ± standard deviation. All data were checked for normality before two-way analyses of variance (two-way ANOVA). All statistical tests with p<0.05 were considered significant. All the transcriptome and metabolome visualizations (Venn diagrams, heat maps, volcano maps, etc.) were made using an online platform (www.majorbio.com). The graphics were drawn using Origin 2021 and Adobe Illustrator CC 2019.

## 3 Results and analysis

### 3.1 Physiological changes and Cr accumulation of *C. indica* under Cr stress

In this study, compared with group Cr0, pH and EC in soil increased significantly after Cr(VI) was added, but their values decreased substantially with increased stress time. Meanwhile, soil organic matter (SOM) content decreased significantly with the increase in stress time. However, it was significantly higher than that of the control group at 14 and 21 days (Figure 1A). In addition, prolonged Cr stress time significantly decreased the biomass of *C. indica*. In addition, with the increased Cr stress time, the contents of carotenoid and total chlorophyll also reduced significantly, especially in group Cr7, which decreased by 50.22% and 33.85% compared with group Cr0 (Figure 1B). Meanwhile, with the increase of Cr stress time, the contents of Cr(VI) and Cr(III) in soil showed a trend of first increasing and then decreasing. In contrast, the Cr content in *C. indica* showed a trend of significantly increasing all the time and reached the maximum value in group Cr21 (Table 1). In addition, it was observed in this study that the biomass and photosynthetic pigment content of *C. indica* showed the maximum value in the CK7 group (Figure 1B, E). In summary, these results indicate that prolonged stress time can significantly inhibit the growth of *C. indica* and increase the contents of Cr of different forms in plants under high Cr stress.

**Fig. 1.**
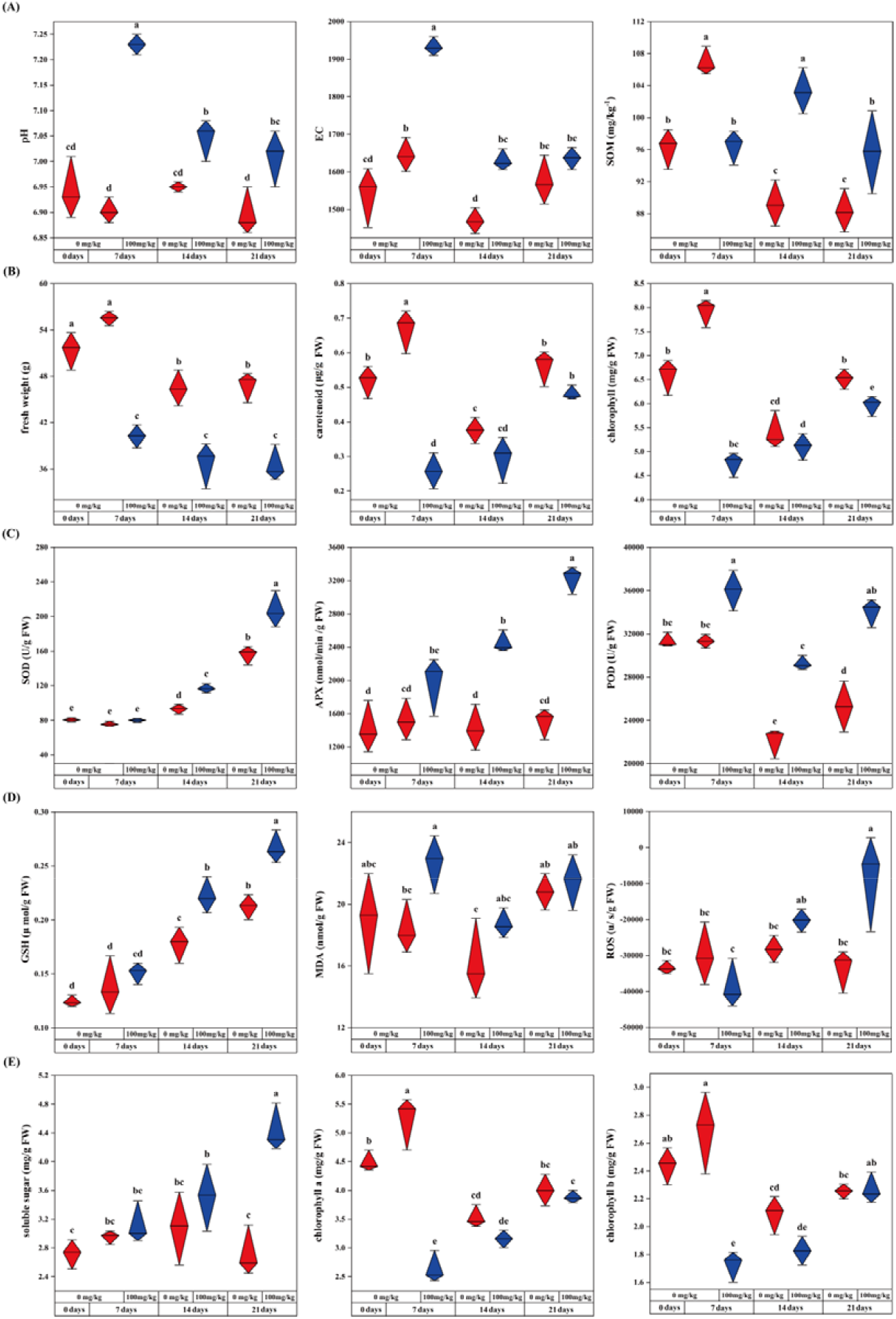
Effects of Cr stress on soil physicochemical properties (pH, EC, and SOM) and physiological and biochemical indexes of C. indica (Freshweight, Carotenoid, Chlorophyll, SOD, APX, POD, GSH, MDA, ROS, Soluble sugar, Chlorophyll a, and Chlorophyll b). Data within the same followed by a string of the same lowercase letters are not significantly different (P > 0.05). At the same time, a series of other letters show a significant difference (P < 0.05).

**Table 1.** Accumulation and transport of heavy metal Cr by *Canna indica*.

| Group | Cr(VI) in soil<br>(mg/kg) | Cr(III) in soil<br>(mg/kg) | Cr(VI) in leaves<br>(mg/kg) | Cr(III) in leaves<br>(mg/kg) |
| --- | --- | --- | --- | --- |
| Cr0 | 12.40±0.26d | 29.36±0.37f | 0.44±0.07d | 1.04±0.11c |
| CK7 | 13.27±0.31b | 37.89±0.20d | 0.39±0.05d | 1.11±0.05c |
| Cr7 | 14.60±0.26a | 117.84±2.05a | 4.75±0.58c | 9.45±0.41b |
| CK14 | 12.30±0.36d | 33.62±0.17e | 0.56±0.05d | 1.19±0.04c |
| Cr14 | 13.10±0.28bc | 85.68±1.32b | 6.22±0.82b | 14.54±0.46a |
| CK21 | 12.50±0.37d | 28.13±0.91f | 0.73±0.06d | 1.36±0.15c |
| Cr21 | 12.67±0.15cd | 65.93±0.96c | 8.82±0.45a | 15.34±1.73a |

Data within the same followed by a string of the same lowercase letters are not significantly different ( $P > 0.05$ ). At the same time, a string of different letters shows a significant difference ( $P < 0.05$ ).

In this study, after the addition of Cr(V), the activities of superoxide dismutase (SOD) and APX(ascorbate peroxidase) in C indica of group Cr21 were increased by 75.49% and 56% compared with that of group Cr0, respectively (Figure 1C). In contrast, POD(peroxidase) activity showed the highest value in the Cr7 group, which increased by 13.02% compared with the Cr0 group. With increased Cr stress time, the activity of antioxidant enzymes showed an increasing trend. In addition, it can be seen from the change of glutathione (GSH) and soluble sugar content in *C. indica* that the content of GSH and soluble sugar is increasing (p<0.05), and its maximum value was found in group Cr21. At the same time, the contents of malondialdehyde (MDA) and reactive oxygen species (ROS) were also significantly increased after Cr(VI) was added, and reached their peaks in Cr7 and Cr21 groups, respectively (Figure 1D, E). These results indicate that Cr stress can interfere with the everyday life activities of plants, induce oxidative stress, and activate the oxidative stress mechanism of plants to cope with Cr stress.

### 3.2 Metabolomic analysis

#### 3.2.1 Metabolic changes of *C. indica* roots under Cr stress

In this study, non-targeted metabolomics (LC-MS) was used to study the metabolism of *C. indica* roots to identify the different metabolites associated with Cr immobilization in *C. indica* roots to understand better the stress response mechanism of *C. indica* roots under Cr stress. Firstly, PCA and PLS-DA analysis were performed on four groups of cluster data with different treatments. PCA analysis could indicate the overall metabolism differences among treatment groups and the degree of variation among each sample. The results showed that Cr stress had little effect on the metabolites in the roots of *C. indica* under cationic mode (Figure 2A). At the same time, there was a significant partitioning phenomenon between Cr14 and Cr21 groups and the Cr0 groups due to the significant differences between groups in the Cion mode (Figure 2A, C).

**Fig. 2.**
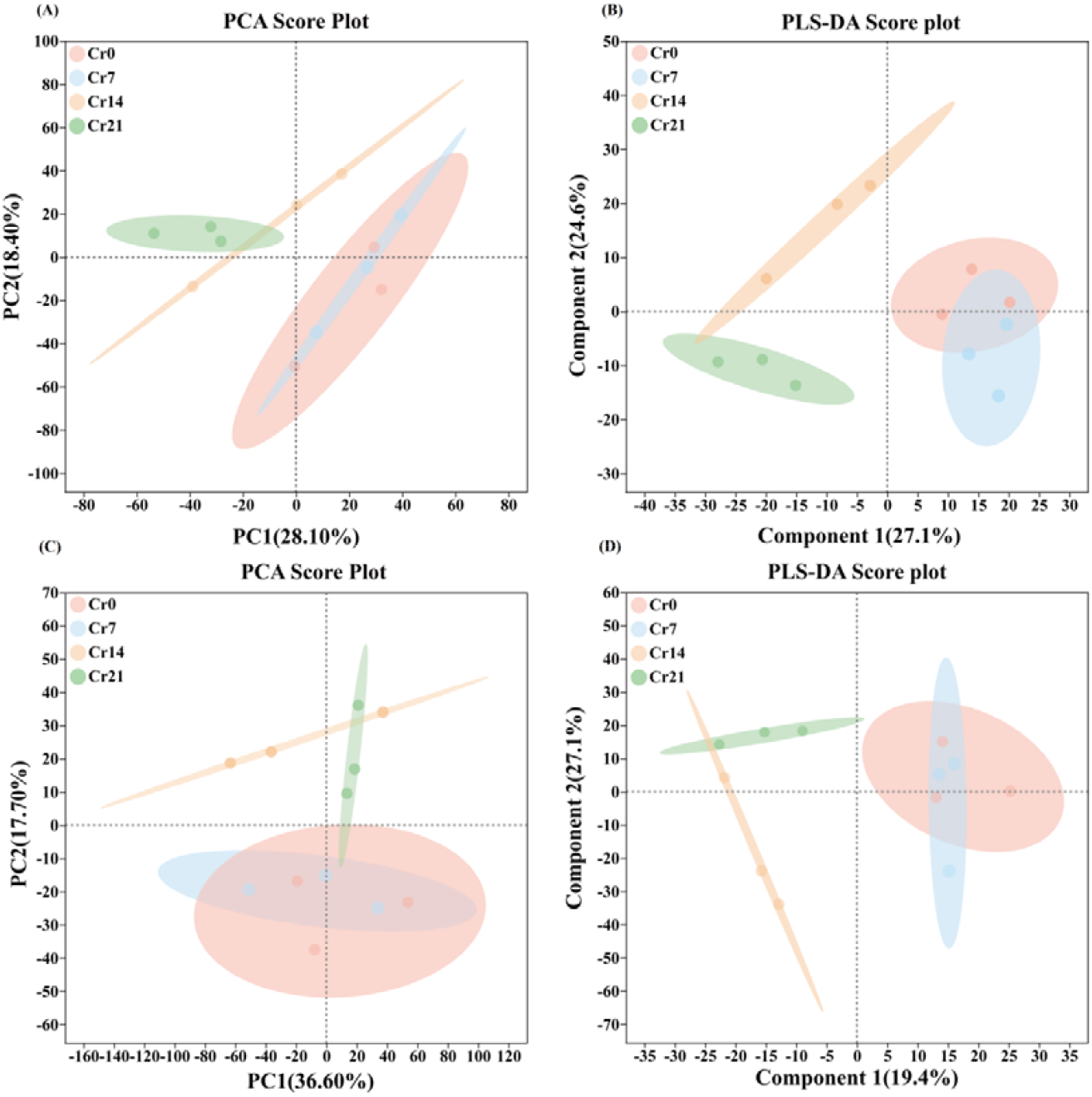
PCA and PLS-DA plot metabolic profiles in C. indica root among different groups in both the positive ion (A, B) and negative ion (C, D) modes under Cr treatments, respectively.

Compared with PCA analysis, PLS-DA analysis is a discriminant analysis method in multivariate data analysis technology, which can effectively distinguish the values between groups and find the variables that affect the differences between groups. This study showed that the PLS-DA scatter in all treatment groups showed an apparent partitioning phenomenon in the cationic mode (Figure 2B, D). Meanwhile, in the cationic mode, the differential interpretation rate of PLS-DA analysis reached 51.7%, which reflected the reliability and applicability of this data for future studies (Figure 2B). In summary, these results suggest that Cr stress can significantly affect the composition of metabolites in the roots of *C. indica*.

#### 3.2.2 Cluster analysis of DEMs in the roots of *C. indica*

In this study, A total of 243 kinds of DEMs were selected by Veen analysis (VIP>1 and P<0.05); three of these were DEMs shared by each comparison group (Figure 3A). At the same time, this study used cluster analysis to study the metabolic changes in the roots of *C. indica*, and showed them in the heat map. In the heat map, red represented up-regulated metabolites, and blue represented down-regulated metabolites. Different color module matrices revealed the differential distribution of metabolites in the roots of *C. indica* under Cr stress, directly expressing the significant differences between the DEMs in the Cr-exposed group and the control group (Figure 3B, C, D). 38 DEMs were detected in the Cr0 vs Cr7 comparison group, with 24 up-regulated and 14 down-regulated metabolites. Up-regulated metabolites include Chrysoidine free base and Citicoline. Similarly, 95 DEMs were detected in the Cr0 vs Cr14 comparison group (15 up-regulated and 80 down-regulated). Gln Val Tyr Asp, Phytosphingosine-1-P, Methyl jasmonate, M-Coumaric acid, and P-Tolualdehyde were particularly abundant in the Cr-contaminated group. As Cr stress duration increased, more up-regulated DEMs were detected in the Cr0 vs Cr21 comparison group. Asp Ile Gln Gly, Gln Val Tyr Asp, L-Aspartic acid, L-Glutamate, M-Coumaric acid, Mevalonic acid, and Flumiclorac-pentyl were most abundant (Figure 3B, C, D; Table S1). Notably, Gentiopicrin was significantly expressed in all three comparison groups (Table S1). Meanwhile, we identified the top ten DEMs in each comparison group based on changes in differential metabolites (Figure S2A). The volcano map visually illustrates DEM changes (Figure S2B).

**Fig. 3.**
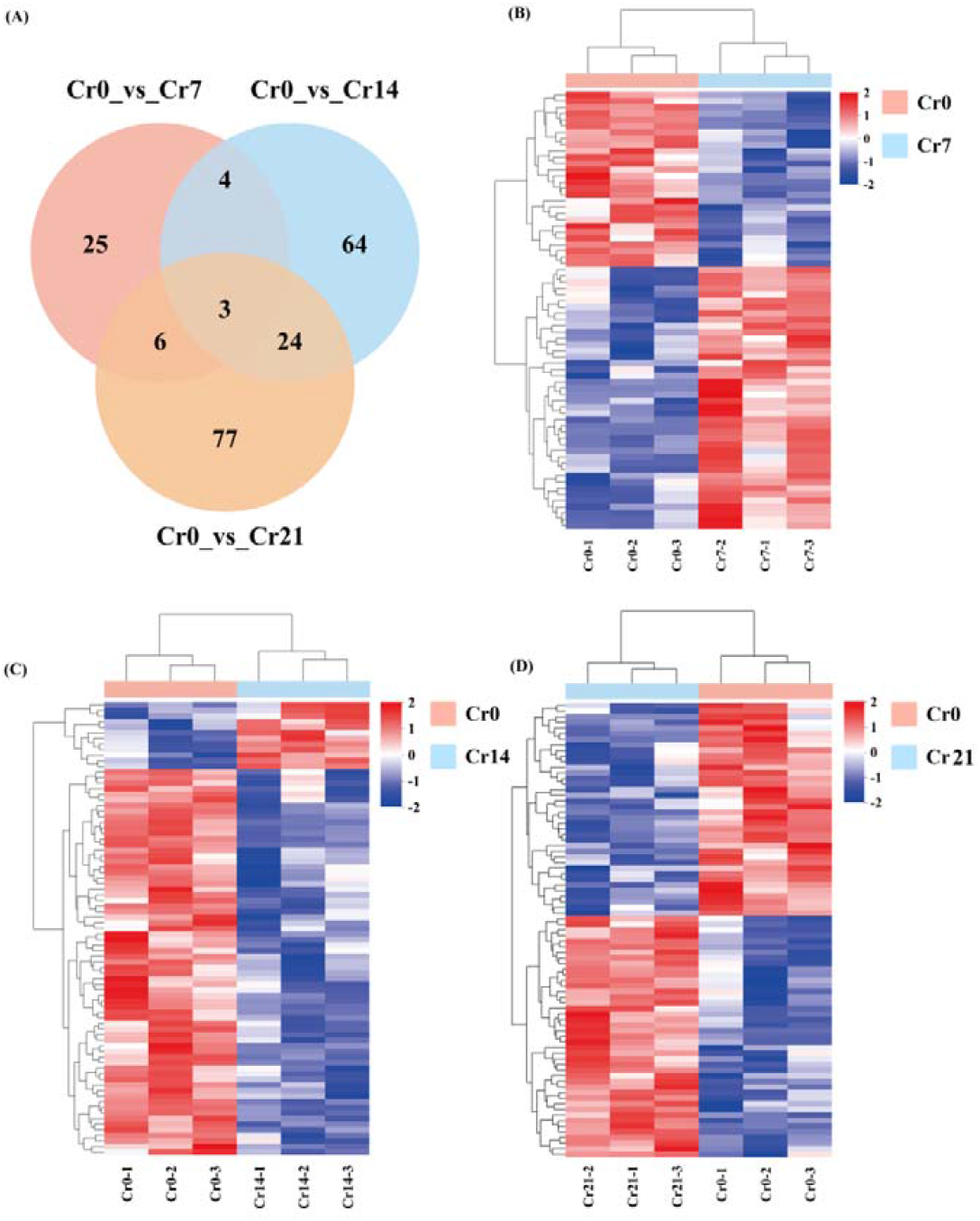
Quality control of Metabolomics data and Changes in DEM expression. (A) Compare the Veen graph of the number of DEMs in groups pairwise. The overlap represents the number of metabolites common to each comparison group, and the non-overlap represents the number of metabolites unique to the comparison group, (B) Heatmap showing the results of the clustering analysis of DEMs

Under Cr stress, Among the secondary metabolite classes, the DEMs are mainly Flavonoids (28.57%), Phenylpropanoids (20%), Terpenoids (31.43%), Fatty acids-related compounds (8.57%), and Alkaloids (5.7%) (Figure S1A; Table S2), in the classification of lipid compounds, It is mainly composed of Fatty acyls (37.5%), Glycerolipids (12.5%), Glycerophospholipids (36.54%), Polyketides (8.65%), and Sterol lipids (8.65%) (Figure S1B; Table S2). These results suggest that amino acids, phenylpropanoids, flavonoids, terpenoids, Fatty acyls, and Glycerophospholipids may play a crucial role in detoxifying Cr in *C. indica* roots.

#### 3.2.3 Analysis of enrichment of KEGG functional pathway in DEMs in *C. indica* roots

Through the enrichment analysis of the KEGG pathway, the biological pathways between different comparison groups were determined to further explore the metabolic mechanism of *C. indica* to Cr stress. First, we searched and annotated the DEMs in *C. indica* roots under different Cr treatments and screened out the top 20 metabolic pathways in enrichment (Figure 4A; Table S3). It mainly includes Purine, Glycerophospholipid, Tyrosine, Arachidonic acid, Phenylalanine, Alanine, aspartate and glutamate metabolism, Phenylpropanoid, Phenylalanine, tyrosine, and tryptophan biosynthesis, ABC transporters. The Biosynthesis of cofactors was mainly enriched in the Cr0 vs Cr7 comparison group. Phenylalanine metabolism, Sphingolipid, N-Glycan, Glycosylphosphatidylinositol (GPI)-anchor biosynthesis, and Autophagy was enriched primarily on the Cr0 vs Cr14 comparison group. With prolonged stress, The significantly enriched metabolic pathways in the Cr0 vs Cr21 comparison group mainly included Histidine, Arachidonic acid, Alanine, aspartate and glutamate, Nicotinate and nicotinamide metabolism, and Arginine biosynthesis. It should be noted that Glycerophospholipid metabolism is significantly enriched in Cr7, Cr14, and Cr21 (Figure 4B, C, D; Table S4). Therefore, these DEMs-enriched pathways in *C. indica* may play an essential role in plant response to Cr stress.

**Fig. 4.**
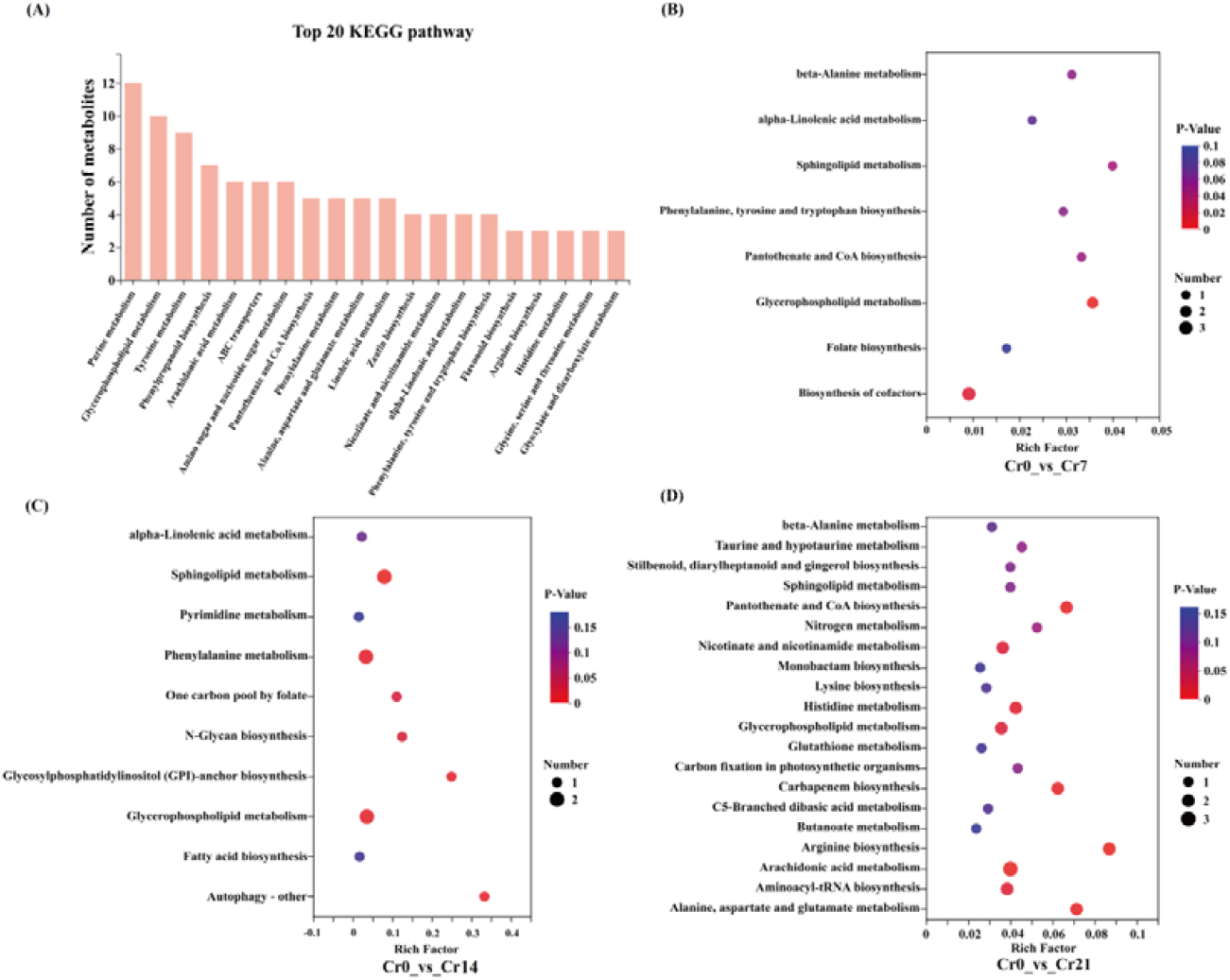
Enrichment analysis of KEGG functional pathways. (A) Top 20 functional pathways for metabolite enrichment. From left to right, the number of metabolites in the column was ranked from high to low. The higher the column, the more metabolites are involved in this pathway among the identified metabolites. (B, C, D) The top 20 pathways of the significance of the up-regulated and down-regulated DEMs on KEGG. The X-axis represented the rich factor, and the Y-axis represented the pathway’s name. The bubble size represents the number of DEMs involved. The bubbles color indicates the enrichment degree of the path.

### 3.3 Transcriptomic analysis

#### 3.3.1 Gene function annotation

Novaseq 6000 (Illumina) was used for transcriptional sequencing to understand further gene expression changes in *C. indica* roots under Cr stress. In this study, 216,527 genes and 516,953 transcripts were identified in 6 different databases, among which the NR database had the highest annotation rate, and 49,065 genes were significantly annotated (Table S5; Figure S3). In addition, a total of 155.92 Gb of Clean Data was obtained from 21 samples in this study, and all Q20 and Q30 values were greater than 98% and 94%, respectively (Table S6). These results reflect the reliability and applicability of this data for future studies.

#### 3.3.2 Analysis of differentially expressed genes (DEGs) and changes in gene expression

In this study, the up-regulated and down-regulated DEGs in the Cr0 group were compared with those in the Cr7, Cr14, and Cr21 treatment groups. The results showed that with the extension of Cr stress time, the up-regulated genes of *C. indica* root increased significantly, with 1393(440 up-regulated and 953 down-regulated) in the three comparison groups, respectively. 6771(2,964 up-regulated and 3,807 down-regulated) and 8083(4,306 up-regulated and 3,777 down-regulated)(Figure 5A). Among them, the top 15 up-regulated and down-regulated genes with differentially expressed levels among the comparison groups are shown in Table S7. Meanwhile, the total number of up-regulated and down-regulated DEGs among the comparison groups was 439 and 133, respectively (Figure 5B, C). In addition, based on the magnitude and significance of the observed stress effects, we further evaluated the overall gene expression between the comparison groups using volcanic maps. We evaluated DEGs in the three comparison groups using volcanic maps (Figure 5D, E, F).

**Fig. 5.**
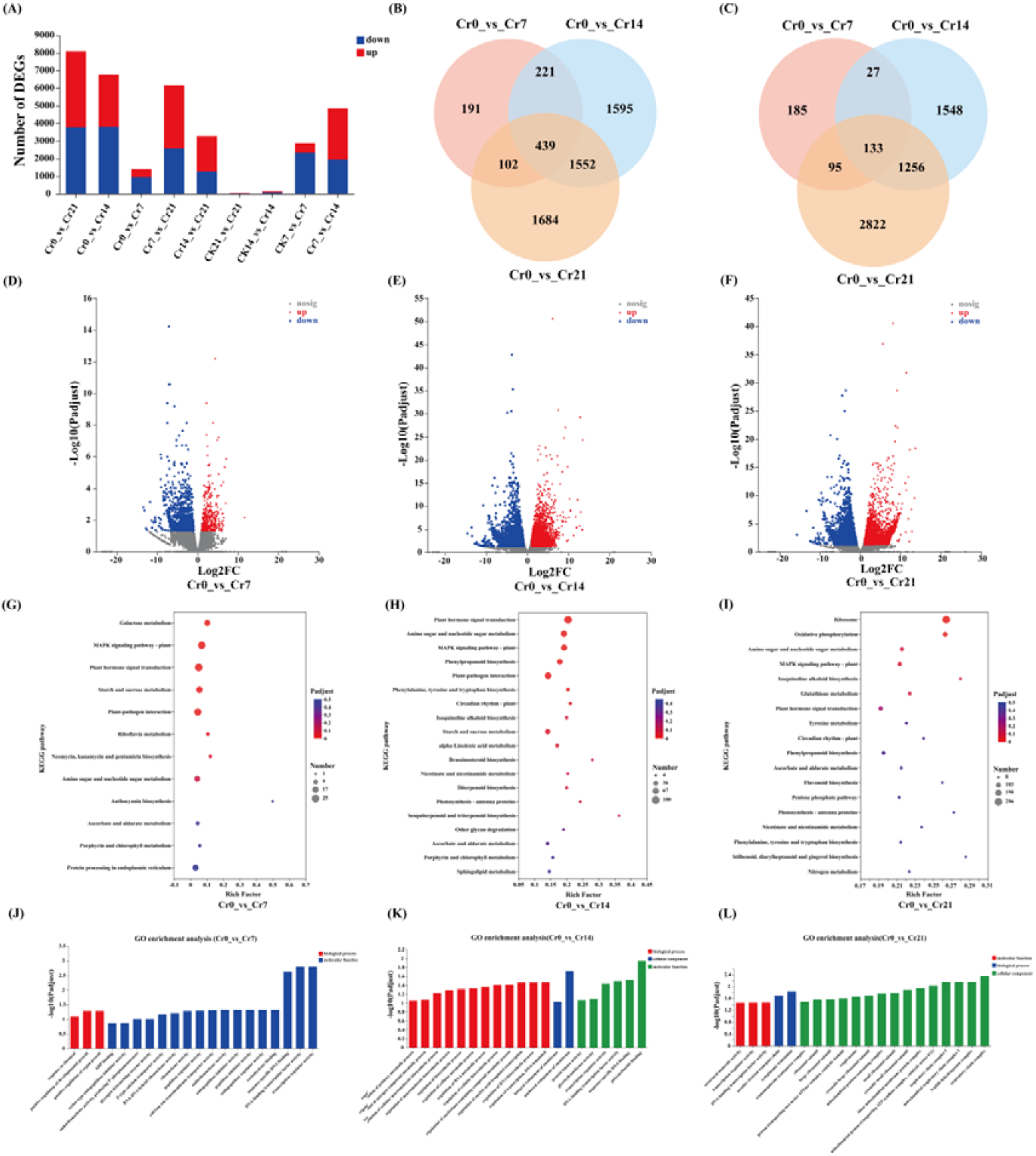
Changes in DEG expression, (A) Up-regulation and down-regulation of DEGs, red represents up-regulation, blue represents down-regulation. (B, C)Veen plot of pairwise comparison of the group’s number of up-regulation and down-regulation DEGs. The overlap means the number of metabolites common to each comparison group, and the non-overlap represents the number of metabolites unique to the comparison group. (D, E, F) Volcano plots of DEGs up-regulation and down-regulation (G, H, I) The top 20 pathways of the significance of the up-regulated and down-regulated DEGs on KEGG. The X-axis represented the rich factor, and the Y-axis represented the pathway’s name. The bubble size represents the number of DEGs involved. The bubbles color indicates the enrichment degree of the path (J, K, L) GO pathway enrichment analysis.

#### 3.3.3 KEGG and GO functional enrichment analysis of DEGs

The biological functions of DEGs in the three comparison groups were elucidated by GO enrichment analysis under Cr stress. In the Cr0 vs Cr7 comparison group, DEGs were significantly enriched in DNA-binding transcription factor activity and transcription regulator activity. Participate in the regulation of RNA biosynthetic process, regulation of transcription, DNA-templated, and an anchored component of membrane DEGs were mainly significantly enriched in the Cr0 vs Cr14 comparison group. With the extension of Cr stress time, The functional pathways significantly enriched in the Cr0 vs Cr21 comparison group mainly include aerobic electron transport chain, cytoplasmic translation, inner mitochondrial membrane protein complex, NADH dehydrogenase complex, and respiratory chain complex (Figure 5J, K, L; Table S9).

KEGG functional pathway enrichment analysis is the primary database for identifying biological pathways between comparison groups. Firstly, the DEGs of *C. indica* roots under different Cr treatments were retrieved and annotated, the top 20 metabolic pathways with the highest enrichment levels were selected, and the enrichment analysis results were displayed with a bubble diagram. In the Cr0 vs Cr7 comparison group, upregulated DEGs are mainly involved in Galactose metabolism, and the DEGs involved in Phenylpropanoid biosynthesis are primarily enriched in the Cr0 vs Cr14 comparison group. With the extension of Cr stress time, The significantly enriched functional pathways in the Cr0 vs Cr21 comparison group mainly included Ribosome, Oxidative phosphorylation, Glutathione metabolism, and Isoquinoline alkaloid biosynthesis. It is worth noting that Starch and sucrose metabolism, MAPK signaling pathway-plant, and Plant hormone signal transduction were significantly enriched in Cr7 and Cr14. However, with the extension of Cr stress time, the enrichment degree of these three functional pathways decreased significantly (Figure 5G, H, I; Table S10). In summary, the results of this study showed that in the early stage of Cr stress, plants mainly respond to Cr stress through the regulation of carbohydrate metabolism, signaling, and transcription factors. With the extension of Cr stress time, Glutathione metabolism, Phenylpropanoid biosynthesis, and oxidative defense system were gradually enhanced in plants.

#### 3.3.4 Analysis of co-expression network of transcription factors and weighted genes

The change of gene expression ultimately regulated the response mechanism of the *C. indica* root system to Cr stress. In this study, a total of 1619 transcription factors (TFs) were identified from 33 transcription factor families. The top 5 families were MYB_superfamily, AP2/ERF, bHLH, C2C2, and NAC (Figure 6A). Weighted gene co-expression network analysis (WGCNA) was used to investigate the relationship between genes and physiology and biochemistry in *C. indica*. Thirty-four gene modules were identified, including 26296 genes (Figure 6B, C). Among them, the central gene of the MEmidnightblue module was positively correlated with the content of photosynthetic pigments (P<0.001), and through the network diagram and correlation heat map, it can be seen that DEGs regulating photosynthetic pigments were significantly up-regulated at the early stage of stress, but significantly down-regulated at the late stage of stress (Figure 6D, E). The central gene of the MEblack module was positively correlated with the degree of lipid peroxidation (MDA) in plants (P<0.001), according to the screened central genes, lipid peroxidation in Cr7 and Cr14 groups was significantly enhanced but significantly weakened at the later stage of stress (Figure 6F, G). The central gene of the MEdarkolivegreen module was positively correlated with the activity of antioxidant enzymes (SOD) and the content of non-enzymatic antioxidant substances (GSH) in plants (P<0.001), whose central gene was significantly enhanced in Cr7 and Cr21 groups (Figure 6H, I). These results suggest that the core gene clusters screened by the gene visualization network in photosynthetic pigment biosynthesis, ROS oxidative stress, and antioxidant mechanisms may play a crucial role in *C. indica*.

**Fig. 6.**
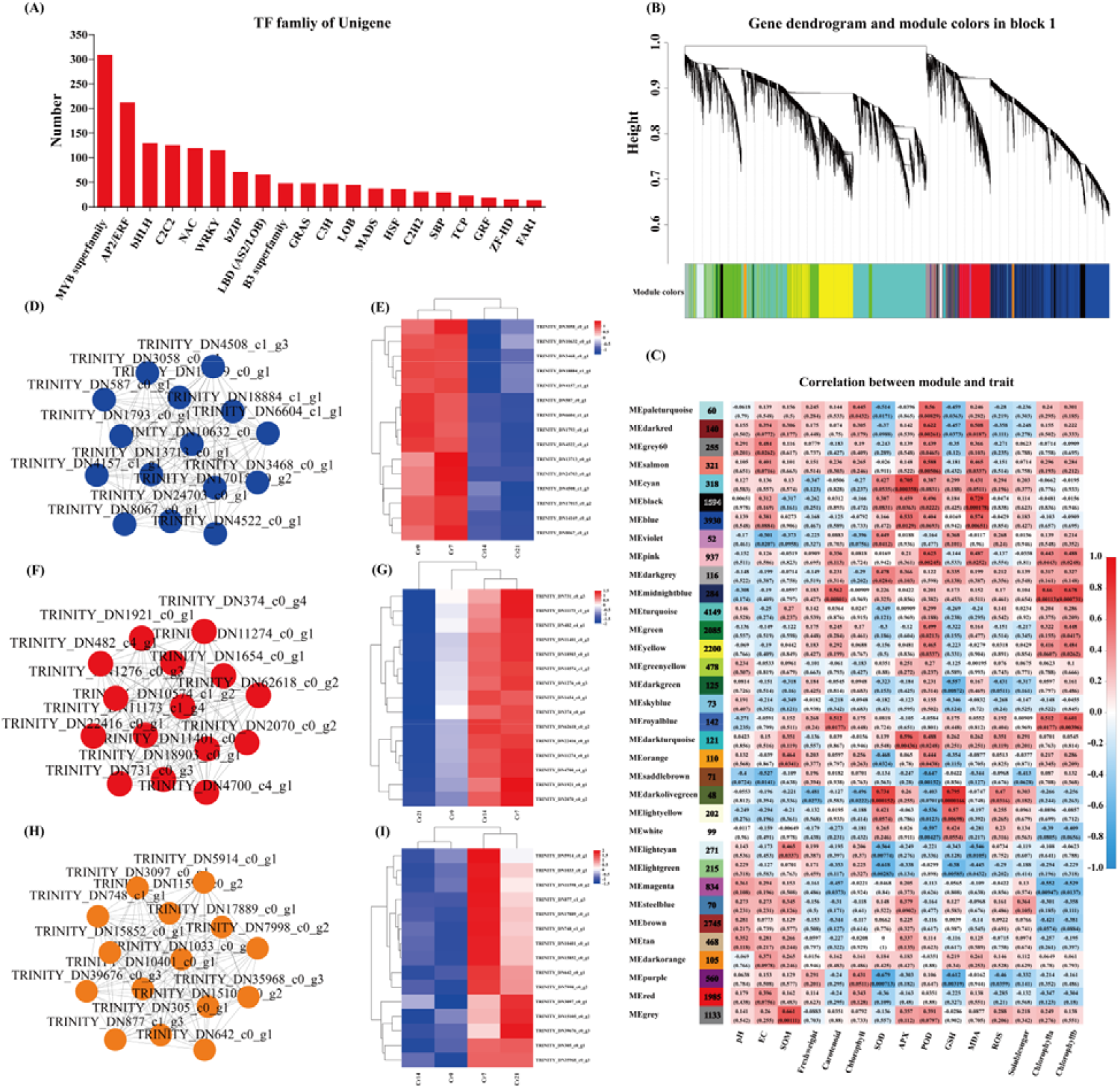
Results of TF and WGCNA. (A) The number of top 20 TFs. (B) Hierarchical clustering tree showing coexpression modules identified by WGCNA. (C) Module sample association relationships. (D, E) midnight blue module central gene symbiosis network map and clustering heat map, (F, G) black module major gene symbiosis network and clustering heat map, (H, I) Symbiotic network map and clustering heat map of dark olive-green module center gene.

## 4 Discussion

### 4.1 Physiological response changes of *C. indica* under Cr stress

Cr is almost not involved in any metabolic pathways in plants, but its toxicity will hinder and affect plant growth and development’s physiological and biochemical processes (Mumtaz et al., 2022). Among them, changes in plant physiological characteristics are usually related to homeostasis and stress strategies (Xu et al., 2022). Plants can enhance their tolerance to heavy metals through their antioxidant mechanisms, energy metabolism, and hormone transduction processes (Pan et al., 2021). This study showed that after *C. indica* was exposed to Cr stress, the contents of carotenoid and chlorophyll in leaves were significantly reduced, and plant growth was significantly inhibited (Figure 1B, E). This is in contrast to previous studies in *Solanum lycopersicum L*.(Anastasis et al., 2021), *rice* (Yu et al., 2018), and *cauliflower* (Ahmad et al., 2020), indicating that plant photosynthesis may be interfered with by Cr stress. Meanwhile, in this study, the central gene screened by WGCNA in the MEmidnightblue module was significantly positively correlated with the content of photosynthetic pigments. However, its gene expression was significantly down-regulated with the extension of stress time (Figure 6D, E), which may explain why photosynthetic pigments decreased during stress. Plants can activate endogenous defense mechanisms in response to ROS-induced oxidative stress. The defense system is mainly composed of antioxidant enzymes and chelates, such as SOD, APX, and POD, which can scavenge free radicals and neutralize intermediates with oxidative toxicity to maintain plants’ homeostasis, thus reducing oxidative damage in plant cells (Chen et al., 2022). In this study, it was observed that the activity of antioxidant enzymes (SOD, APX, and POD) and the contents of GSH and soluble sugar increased significantly with the extension of Cr stress time, which was consistent with the results of previous studies. Among them, soluble sugar could not only serve as energy storage substances for plants but also as signal transduction and osmotic regulation substances, playing a pivotal role in plant growth and development and stress response (Wei et al., 2019). GSH is a widely recognized essential plant metabolite and functions as an antioxidant and detoxifier, a precursor to phytochelatin. It can bind with Cr, Pb, and other heavy metals, significantly reduce its mobility and bioavailability, and greatly enhance plant tolerance to heavy metals (Mumtaz et al., 2022; Meng et al., 2022). It should be noted that the oxidative stress system of *C. indica* does not activate to the highest level at the initial stage of stress but gradually strengthens with the extension of stress time.

### 4.2 Transcriptional metabolic response of *C. indica* to Cr stress under time series

In Cr stressed environment, plant roots can jointly resist heavy metal poisoning by changing the content of their metabolites and gene expression of critical metabolic pathways (Wang et al., 2022). This study revealed the molecular response mechanism of *C. indica* under Cr stress under time series through untargeted metabonomics studies combined with transcriptomic analysis. The results showed that DEGs involved in Galactose, Starch, and sucrose metabolism were significantly up-regulated in *C. indica* under short-term Cr stress, leading to a significant increase in carbohydrate content in *C. indica* (Figure 7A, B; Figure 3B). Many studies have shown that the upregulation of endoglucanase, EGLC, SUS, and β-amylase in the Starch and sucrose metabolism pathway will increase soluble sugar content in plants (Meng et al., 2022). This phenomenon is essential in plant growth, development, and signal transduction in response to Cr stress (Wang et al., 2021c). In addition, galactose metabolism is an essential intermediate process of the carbohydrate cycle in plants, providing precursors for glucose metabolism and energy support for metabolic processes under stress, thus helping plants maintain nutritional balance in harsh environments (Yu et al., 2023). Therefore, in the early stage of Cr stress, the rich carbohydrate metabolism process in *C. indica* may help to enhance plant tolerance to Cr for a short period.

**Fig. 7.**
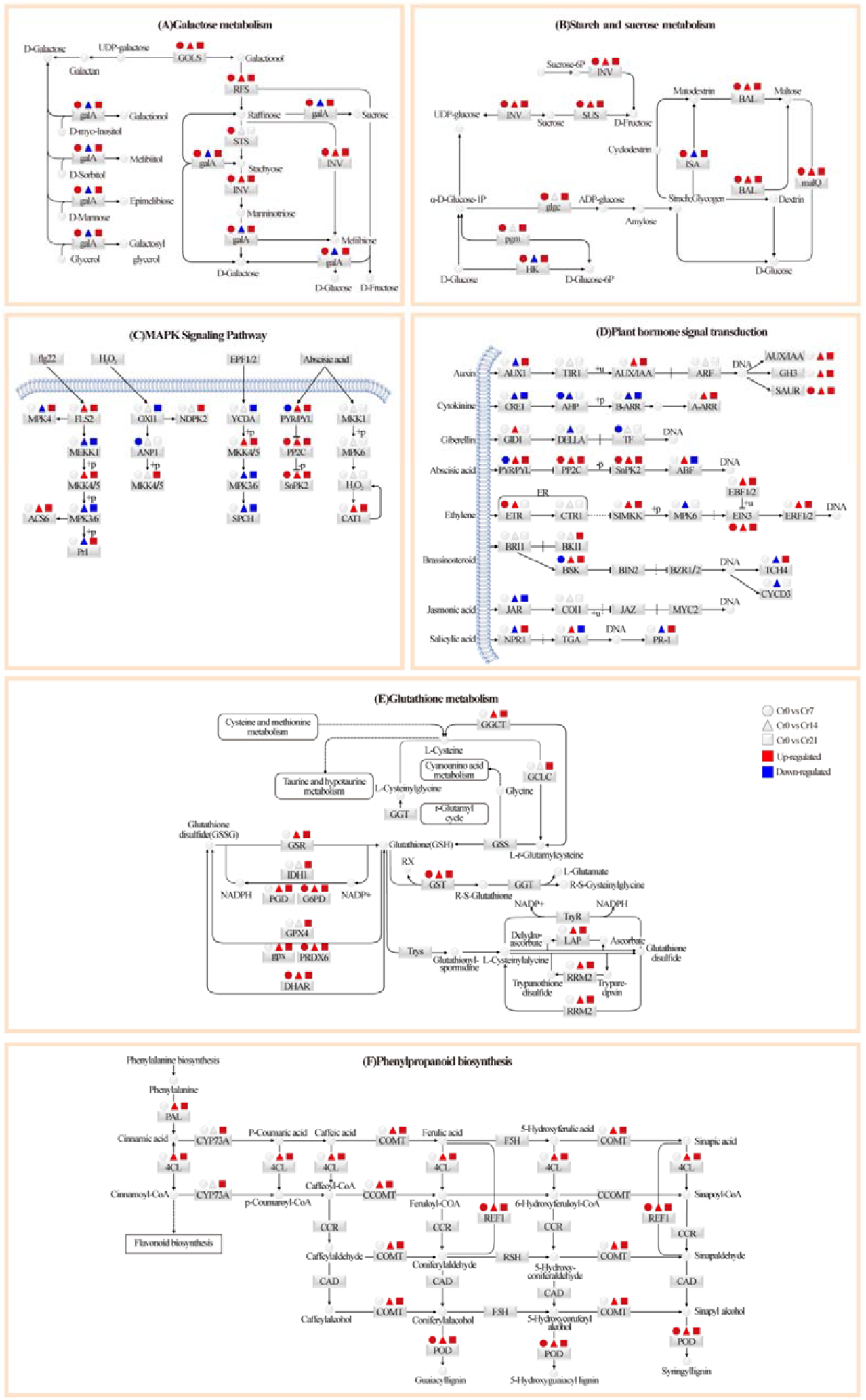
Changes of main metabolic pathways in C. indica after Cr stress. Based on the data generated by the KEGG database and with some modifications, the path was established. Red and blue indicate up-regulation and down-regulation, respectively, while gray indicates no significant change.

In this study, in the middle stage of Cr stress, it was also found that plant hormone signal transduction and MAPK signaling pathway were significantly enhanced in *C. indica* (Figure 7C, D). Generally speaking, plant hormones are vital regulatory factors mediating stress response, which can rapidly activate stress response mechanisms in various organelles, thus reducing oxidative damage in plants (Meng et al., 2022; Mittler et al., 2022). In order to induce the expression of various transport factors and the production of the Cr(VI) detoxification peptide chain, Enzymes involved in metabolic pathways in plants are activated in response to stress signals (Mumtaz et al., 2022). Heavy metals promote the expression of TFs and stress-responsive genes in plants and activate various signaling pathways, including MAPK signaling and hormone signaling (Kumar et al., 2020; Star et al., 2019). In existing studies, Kumar et al. (2015) used *Arabidopsis thaliana* to reveal the activation pathway of MAPK signaling pathway in plants under heavy metal stress, mainly including ROS accumulation and changes in the antioxidant system. This study found that flg22 and Abscisic acid-related genes in the MAPK signaling pathway were significantly up-regulated in *C. indica* after the exogenous addition of Cr(VI) at the middle stage of stress. In auxin signal transduction, upregulation of ARF, which is a hub gene, in a high-Cr environment not only regulates the upregulation of downstream SAUR and GH3 to cope with adverse environmental conditions but also regulates the direction of plant growth factors by binding to Aux/IAA repressor proteins (Figure 7C; Guilfoyle et al., 2007). This indicates that under Cr stress, with the extension of time, the growth strategy of *C. indica* changes, and the rapid root growth is conducive to the absorption of soil nutrients by plants, thus conducive to the survival of plants in the stressed environment. Studies have shown that ABA is closely related to the signal transduction of plant stress resistance (Cutler et al., 2010). The results showed that in the middle stage of Cr stress, ABA receptor PYR/PYL-related genes were up-regulated and inhibited downstream PP2C and SnPK2-related genes, thus activating ABF binding factors and enhancing the ABA signal transduction process. In the JA signaling pathway, Verma et al. (2020) used transcriptomics to reveal the signal regulation of MYC2 in Arabidopsis thaliana, and the results showed that MYC2 mainly mediated the biosynthesis of proline. Meanwhile, in the salicylic acid signaling pathway, the results of Zhang et al. (2022b) showed that the NPR1 protein involved in REDOX regulation was mainly mediated by salicylic acid. Therefore, we speculate that IAA, ABA, jasmonic acid, and salicylic acid play a crucial role in enhancing Cr tolerance in plants.

The plant can regulate heavy metals’ transport through cell wall fixation, metal chelation, and vacuolar compartments. Among them, the absorption of heavy metals is mainly concentrated in the plant root cell wall, which can prevent external pollutants from entering the cell interior (Yuan et al., 2022a). Solanum nigrum L.(Wang et al., 2022), *Celosia argentea Linn* (Yu et al., 2023), and *Sedum alfredii* (Ge et al., 2023), Studies on the distribution of heavy metals that the root cell wall is the leading site for binding heavy metals and plays a crucial role in plant response to Cr stress. The present study showed that after the exogenous addition of Cr(VI), DEGs involved in the regulation of phenylpropanoid biosynthesis was significantly up-regulated at the later stage of stress with the extension of stress time (Figure 5H, I; Figure 7E). Therefore, we speculated that *C. indica*, under long-term Cr stress, may activate the phenylpropanoid biosynthesis pathway in plants due to the increased accumulation of HMC in the roots, affecting the synthesis of coumarin and lignans, thus reducing the mobility and bioavailability of Cr. This can enhance the tolerance of C. indica to HMC (Sharma et al., 2020; Xian et al., 2020). Wang et al. (2021b) also showed in recent studies that plants under heavy metal stress could enhance their binding ability with heavy metals by inducing their cell wall metabolism and reshaping their structure to enhance their tolerance to heavy metals. In addition, plants can jointly regulate the accumulation and transport of heavy metals by increasing cellulose and pectin contents and xylem cell wall thickness to alleviate heavy metals’ toxic effects on plants (Guo et al., 2021a). Phenylpropanes (Figure S1) and amino acid (Phenylalanine) contents (Figure 3C, D) and the Phenylalanine metabolism pathway (Figure 4C, D) were significantly up-regulated in late Cr stress. Among them, phenylalanine is catalyzed and oxidized to tyrosine by phenylalanine hydroxylase, which, together with tyrosine, synthesizes important hormone substances and participates in glucose metabolism and lipid metabolism (Adams et al., 2019). Moreover, it can be converted into phenylpropane metabolites in secondary metabolic biosynthesis, including lignin and flavonoids, which greatly enhance the tolerance of plants to heavy metals (Yuan et al., 2022a). In addition, the enhancement of the Phenylalanine metabolism pathway is consistent with transcriptome results, suggesting that cell wall metabolism-related pathways of *C. indica* may play a crucial role in plant response to heavy metal stress under prolonged heavy metal stress (Yuan et al., 2022a).

GSH is a plant antioxidant, mainly in the form of reduced glutathione (GSH) and oxidized glutathione (GSSG) in plants. It is involved primarily in the removal of ROS in plants and the chelation of heavy metals and plays an essential role in the stress resistance of plant cells (Yu et al., 2023). This study showed that DEGs involved in regulating GSH metabolism in *C. indica* roots were significantly up-regulated under two treatment times of Cr14 and Cr21 (Figure 5H, I; Figure 7F). This may be because plants exposed to heavy metal stress for a long time can produce a large amount of ROS. However, excessive ROS accumulation will eventually lead to lipid peroxidation of the cell membrane and damage its function of the cell membrane. Therefore, genes that regulate GSH metabolism in plants are induced to be expressed in response to oxidative damage caused by stress (Mumtaz et al., 2022). However, some studies have shown that ROS in plants can be independently produced in different compartments and serve stress-sensing and signaling purposes in plants to regulate gene expression and induce stress recovery (Mittler et al., 2022). It is worth noting that GSH can not only capture and bind heavy metal ions attached to the enzyme protein sulfhydryl but also reduce them to acidic substances through the combination of sulfhydryl with free radicals in plants to accelerate the removal of free radicals, thus enhancing the tolerance of plants to heavy metals (Li et al., 2021). Meanwhile, Yu et al. (2023) revealed the detoxification mechanism of GSH in plants of *Celosia argentea Linn* under heavy metal stress through multi-omics analysis. The results showed that GSH is a precursor to phytochelatin peptides (PCs) using a synthase to complex free HM ions in plant cells. This HM complex is then delivered to plant vacuoles via tonoplast membrane transporters for eventual detoxification. Hasanuzzaman et al. (2020) reported that plants’ antioxidant content would increase significantly under oxidative stress, dramatically enhancing plants’ stress resistance. In addition, WGCNA showed that the central gene of the MEdarkolivegreen module was significantly positively correlated with the GSH content in plants (P<0.001), and its expression was significantly increased in the Cr21 group (Figure 6H, I). This is similar to the gene expression results of the GSH metabolic pathway in Morus alba L. and Pepper (Guo et al., 2021b; Mumtaz et al., 2022). Key functional genes involved in glutathione S-transferase and Glutathione metabolism were significantly up-regulated under heavy metal stress, which greatly enhanced plant stress resistance.

The contents of amino acids (L-Glutamate, Glutamine, and Glycine) and flavonoid metabolites at the late stage of Cr stress (Figure 3D; Figure S1) and glutamate metabolism (Figure 4D) were significantly up-regulated. In plants, amino acids can effectively chelate metal ions in the cytoplasm to reduce the toxic effect of heavy metals on plants (Singh et al., 2016). L-Glutamate is an intermediate for the biosynthesis of oxidized amino acids, which plants can use for secondary metabolisms, such as the biosynthesis of glutamine, proline, and lysine (Yuan et al., 2022b). Glutamine is an essential precursor of glutathione biosynthesis in plants. Glycine is an antioxidant involved in heavy metal chelation in plants, and the presence of these metabolites plays a crucial role in plant survival and stress resistance (Feng et al., 2021). In addition, proline is not only an ideal osmotic regulator in plants but also a protective substance for enzymes in cell membranes and plants, as well as a free radical scavenger (Meng et al., 2022), which can protect the growth of *C. indica* under heavy metal stress. Mwamba et al. (2022) revealed the adaptive mechanism of *Brassicanapus* to heavy metal stress through metabonomics analysis. The results showed that phenolic compounds were involved in response to heavy metal stress, among which flavonoids were significantly induced, and indicated that the antioxidant mechanism was the final response strategy to heavy metal stress. In conclusion, *C. indica* can resist Cr stress mainly through carbohydrate metabolism (galactose, Starch, and sucrose metabolism) in the early stage of Cr stress (Cr7). With the extension of Cr stress time, in the middle stage of stress (Cr14), plants mainly use plant hormone signal transduction and MAPK signaling pathway to regulate their stress response system to cope with Cr stress jointly. In anaphase of stress (Cr21), *C. indica* mainly activates its Glutathione metabolism and Phenylpropanoid The expression of genes related to biosynthesis and the accumulation of metabolites (phenylpropanoids, phenylalanine, flavonoids, and L-Glutamate) can jointly resist the toxicity of Cr through endogenous and exogenous molecular response defense mechanisms, thus enhancing the tolerance of plants to Cr stress.

## 5. Conclusions

This study showed that *C. indica* showed corresponding changes in plant growth and physiological metabolite levels under Cr stress at different exposure times. Due to the toxicity of Cr itself, plant growth was significantly inhibited, the biosynthesis of photosynthetic pigments was damaged, and the expression levels of enzymes were promoted. Non-enzymes-promoted antioxidants in plants were reduced considerably. In addition, this study revealed the molecular response mechanism of *C. indica* under different stress times through transcriptome and metabolomics analysis. The results showed that *C. indica* maintained its energy supply through Galactose, Starch, and sucrose metabolism in the early stages of stress (Cr7), ensuring the regular operation of liver metabolism and resisting the toxicity of Cr. With the extension of stress time, the plant hormone signal transduction and MAPK signaling pathway processes of *C. indica* during the middle stage of stress (Cr14) are affected, thus altering the plant growth strategy. Finally, in the later stages of stress (Cr21), *C. indica* mainly activates the expression of Glutathione metabolism and Phenylpropanoid biosynthesis genes and the accumulation of antioxidant substances (phenylalanine, flavonoids, and L-Glutamate). Both endogenous and exogenous defense mechanisms can jointly resist Cr poisoning, enhancing the tolerance of *C. indica* to Cr. The results of this study are expected to provide new insights into the growth, physiological and molecular response mechanisms of wetland plants under Cr stress.

## Acknowledgments

This study is financially supported by the National Natural Science Foundation of China (31560107), and by the Science and Technology Support Project of Guizhou province, China (Guizhou Branch Support [2018]2807).

